# Attractors are less stable than their basins: Canalization creates a coherence gap in gene regulatory networks

**DOI:** 10.1101/2025.11.06.687062

**Authors:** Venkata Sai Narayana Bavisetty, Matthew Wheeler, Claus Kadelka

## Abstract

Waddington’s epigenetic landscape has served as biology’s central metaphor for cellular differentiation for over half a century, depicting mature cell types as balls resting in stable valley floors. Boolean networks – introduced by Kauffman in 1969 to model gene regulatory dynamics – provide a mathematical formalization of this landscape, where attractors represent phenotypes and basins of attraction correspond to developmental valleys. Traditional stability measures quantify robustness by perturbing arbitrary states, yet biological systems typically reside at attractors rather than in transient states. Here we formalize and systematically analyze attractor coherence – a stability measure Kauffman originally envisioned but never rigorously developed – which quantifies how likely a perturbation of an attractor state causes phenotype switching. Analyzing 122 expert-curated biological Boolean models, we reveal a striking paradox: attractors representing mature cell types are consistently less stable than the developmental trajectories approaching them. Largescale simulations of random networks demonstrate that this coherence gap arises from canalization – a hallmark of biological regulation where individual genes can override others. While canalization increases overall network stability, it disproportionately stabilizes transient states, positioning attractors near basin boundaries. The gap’s magnitude is almost perfectly predicted by network bias (Spearman’s *ρ* = -0.997), itself modulated by canalization. These findings revise Waddington’s landscape: canalization carves deep protective valleys ensuring developmental robustness, yet simultaneously flattens ridges near valley floors, facilitating phenotypic plasticity when multiple fates coexist. This explains how biological systems achieve both reliable development and plasticity, with implications for understanding development, disease-related transitions, and designing robust yet controllable synthetic gene circuits.

## Introduction

Waddington’s epigenetic landscape metaphor has profoundly shaped how biologists conceptualize cellular differentiation and stability [1, 2, 3]. In this metaphor, a ball rolling down a landscape represents a cell progressing through development, with valleys corresponding to different cell fates or phenotypes. The depth of these valleys represents stability: once a cell reaches the valley floor – a mature, differentiated state – it should be difficult to perturb into a different phenotype. This intuitive picture suggests that mature cell types occupy the most stable positions in the regulatory landscape, protected by steep barriers from phenotype-switching perturbations.

Boolean networks (BNs) provide a natural mathematical formalization of Waddington’s landscape [4]. In this framework, genes and proteins are represented by binary nodes, regulatory interactions by edges, and the dynamics evolve according to discrete update rules. The phenotypes or cell types correspond to *attractors* – fixed points or limit cycles to which the system eventually converges. The basin of attraction surrounding each attractor comprises all states that eventually transition to it, directly analogous to valleys in Waddington’s landscape. This mapping has made Boolean networks a powerful tool for studying gene regulatory dynamics across diverse biological processes, from developmental programs to cancer signaling pathways [5].

The stability of Boolean network attractors has been a central question since Kauffman’s foundational work on metabolic stability and epigenesis [4]. Early metrics like the Derrida value [6] characterized dynamical regimes – ordered, chaotic, or critical – by measuring sensitivity to perturbations after a single update. These analyses led to the criticality hypothesis: that biological systems operate near the critical regime to optimize the trade-off between robustness and adaptability [7, 8, 9, 10]. However, the Derrida value fails to capture the influence of network topology and feedback loops [11], depending only on local sensitivities rather than global structure [12, 13]. To address these limitations, *coherence* was introduced as a measure of phenotypic robustness [14, 15, 16, 17]. Coherence quantifies the probability that a single-node perturbation does not cause a transition to a different basin of attraction. Importantly, coherence is computed as an average over all states in the state space – both transient states along developmental trajectories and attractor states representing mature phenotypes. Yet biological systems predominantly reside at attractors, not in transient states. A cell spends the vast majority of its lifetime in a differentiated state, not in the process of differentiating. This raises a fundamental question: Are attractors themselves as stable as the basins surrounding them?

Kauffman himself recognized this distinction in his original 1969 paper, informally introducing the concept of attractor-specific stability [4]. However, this concept was never formalized or systematically analyzed. Here, we formalize *attractor coherence* – the stability of attractor states themselves to perturbations – and distinguish it from *basin coherence*, which measures the average stability of all states leading to an attractor. By comparing these two measures across 122 expert-curated biological Boolean network models and large ensembles of random networks, we reveal a striking and counterintuitive pattern: attractors are often less stable than the developmental trajectories that lead to them.

Large-scale simulations of random networks demonstrate that this coherence gap is not incidental but arises from fundamental design principles of biological regulation. Canalization – the property whereby individual regulatory inputs can override others – is a hallmark of biological gene regulatory networks [18, 8, 10]. We show that while canalization increases overall network stability, it increases basin coherence more strongly than attractor coherence, creating a stability gradient within basins. Attractors become positioned closer to basin boundaries, making them more vulnerable to phenotype-switching perturbations. The magnitude of this coherence gap is almost perfectly predicted by network bias, which is itself strongly modulated by canalization [12, 19].

These findings revise Waddington’s landscape metaphor (Fig. 1). While canalization creates deep valleys that robustly guide cells along developmental trajectories – explaining why development is highly reproducible – it also creates a stability gradient within valleys. Attractors reside in regions with relatively flatter boundaries compared to mid-trajectory states, facilitating phenotype switching when multiple fates are accessible. This architectural feature is most biologically relevant when multiple phenotypically meaningful attractors coexist. It helps explain how biological systems achieve both developmental robustness (high absolute coherence protects trajectories) and phenotypic plasticity (coherence gap enables regulated transitions), with important implications for understanding processes like stem cell reprogramming [20], wound healing, and disease-related transitions such as cancer differentiation [2, 21, 22].

**Figure 1:**
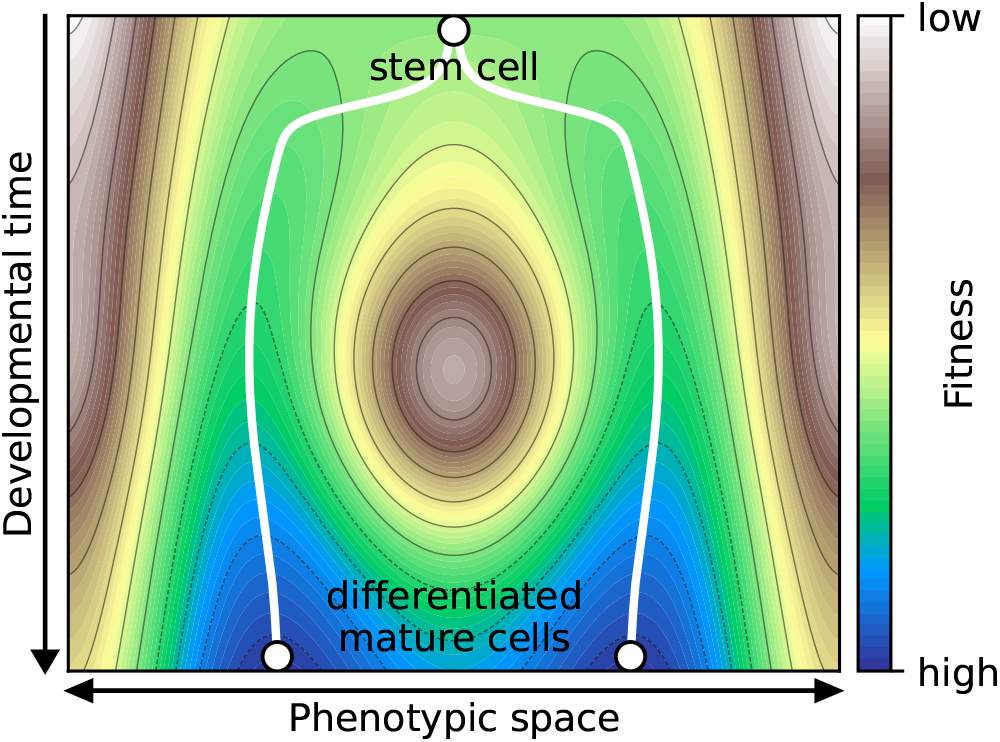
Canalization creates a coherence gap in Waddington’s epigenetic landscape. While canalization increases both basin and attractor coherence in absolute terms, it increases basin coherence more, creating a stability gradient within each valley. Attractors (representing mature cell types or phenotypes) are positioned closer to basin boundaries relative to transient states in their developmental trajectories. The ridges between valleys become relatively flatter near attractors compared to mid-trajectory, making phenotype switching more likely from mature states when multiple phenotypes are accessible. This coherence gap is almost perfectly predicted by network bias, which is itself modulated by canalization.

## Methods

### Boolean Networks

A *Boolean network* (BN) is a finite dynamical system, given by a function

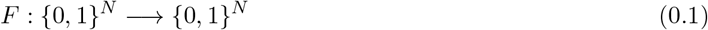

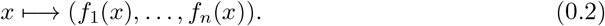

It consists of *N* nodes, where each node can take on only two values, 0 (e.g., OFF, unexpressed, absent) and 1 (e.g., ON, expressed, present). The state of the entire BN at time *t* is represented by the vector **x**(*t*) = (*x*_1_(*t*), …, *x*_*N*_ (*t*)) ∈ {0, 1}^*N*^ . The *state space* {0, 1}^*N*^ is the set of all possible states and coincides with the vertices of the *N* -dimensional Boolean hypercube. Every node is updated in discrete time steps. The state of node *i* at time *t* + 1 is determined by its *update function f*_*i*_ :

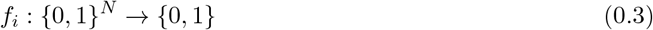

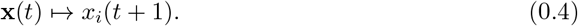

In a given BN, *f*_*i*_ may depend only on a subset of nodes. The *in-degree* of node *i*, is the number of essential inputs of *f*_*i*_.

In this paper, we focus on synchronously updated BNs, where all nodes are updated simultaneously, yielding deterministic dynamics:

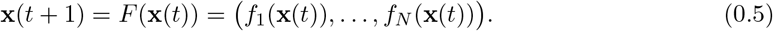

The dynamics are fully described by the *state transition graph*, whose vertices are the 2^*N*^ possible states **x** = (*x*_1_, …, *x*_*N*_ ) ∈ {0, 1}^*N*^ and there is an edge from state **x** to state **y** if *F* (**x**) = **y**. Because BNs have finite size and deterministic dynamics, they eventually exhibit periodic behavior. That is, they settle into *attractors*, which can be fixed points (also known as steady states) or limit cycles. In biological contexts, attractors represent phenotypes or cell types.

Given a BN *F*, let 𝒜(*F* ) denote the set of all its attractors. The *basin of attraction B*_*F*_ (*A*) of a given attractor *A* ∈ 𝒜(*F* ) consists of all states that eventually transition to *A*. Consider the function

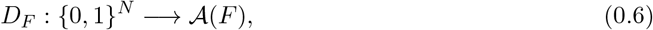

which assigns each state to its attractor. With this notation, we have

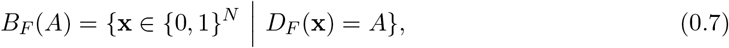

so that

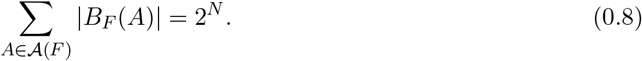

### Coherence

Coherence quantifies the probability that a single-node perturbation does not result in a change of phenotype [14]. Given a state **x** of a BN *F*, let **x** ⊕ *e*_*i*_ denote the state where the *i*-th bit in **x** is flipped. The *coherence* of state **x**, denoted *ψ*_**x**_, is the fraction of neighboring states **x** ⊕*e*_1_, …, **x** ⊕*e*_*n*_ that are in the same basin of attraction as **x**. That is,

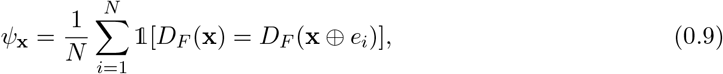

where 𝟙[·] is the indicator function. The *basin coherence ψ*_*B*_ is defined as the average coherence of the states within a basin of attraction *B*, and is given by

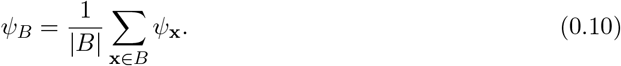

Similarly, the network coherence *ψ*_*F*_ is given by

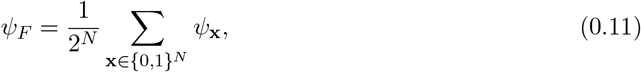

which is simply the sum of all basin coherences, weighted by the relative basin sizes.

### Attractor coherence

Conventional stability metrics for BNs (e.g., the coherence, fragility, or Derrida values) describe the effect of perturbing a random network state (see e.g. [16]). Since most BNs possess many more transient states than attractor states, these stability measures primarily reflect the resilience of transient states to perturbations rather than that of the attractors. However, an unperturbed dynamical system rests at an attractor state. Therefore, to accurately describe the resilience of realworld systems, it is necessary to use a measure that quantifies the stability of attractors themselves. Motivated by this, we study *attractor coherence*, a measure closely related to the previously defined basin coherence. It was already informally introduced by Kauffman in 1969 [4] (also see [23]). Attractor coherence describes the average coherence of states within an attractor, measuring the probability that a single-node perturbation of an attractor state does not push the system into another attractor. Analogous to basin coherence, the coherence of an attractor *A* ∈ 𝒜(*F* ) can be defined by

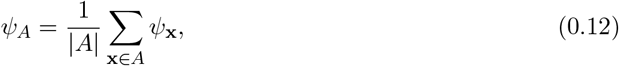

where |*A*| denotes the periodicity of the attractor. If the average coherence of transient states and attractor states differs, this has implications for our understanding of the dynamical robustness of BNs. As an example, consider the 3-node BNs

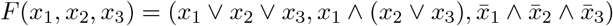

and

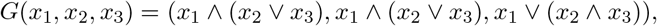

whose state transition graphs are shown in Fig. 2. Both networks have two steady state attractors with same basins of sizes 5 and 3. The basin coherences are 2*/*3 and 4*/*9, respectively, which represent the maximum possible coherence for basins of these sizes [24]. The total network coherence is thus 5*/*8 · 2*/*3 + 3*/*8 · 4*/*9 = 14*/*24. However, the attractor coherences between the two networks differ strongly. *F* has steady state attractors 100 and 110, which both have coherence 1*/*3 because only one of each of their three neighboring states is in their basin of attraction. On the contrary, the steady state attractors 000 and 111 of *G* have much higher attractor coherence of 1 and 2*/*3, respectively.

**Figure 2:**
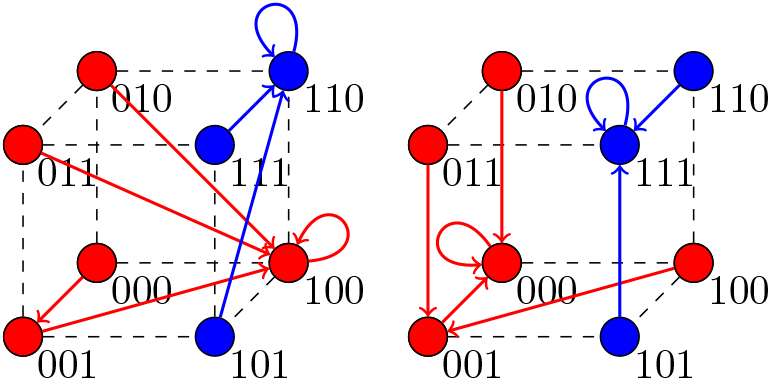
State transition graph of two 3-node Boolean networks. Despite identical basin sizes and basin coherence, both attractors in the left network *F* have coherence 1/3, whereas the attractors 000 and 111 in the right network *G* have coherence 1 and 2/3, respectively. This demonstrates that attractor coherence is not determined solely by basin properties.

### Coherence gap

Our main goal is to understand how the average basin coherence *ψ*_*B*_, the attractor coherence *ψ*_*A*_, and the difference *ψ*_*B*_ − *ψ*_*A*_ vary between network ensembles. The expected values depend on the distribution of relative basin sizes, *ϕ*(*s*). Specifically, we have

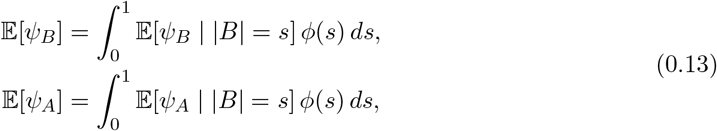

which shows that the overall expectations are weighted by the frequency of basin sizes, which tends to be skewed towards both very small and very large basins.

Different ensembles of BNs may possess different basin size distributions. For example, it is known that the number of attractors increases as the sensitivity of a network to perturbations increases [25]. To enable a comparison of *ψ*_*B*_, *ψ*_*A*_, and *ψ*_*B*_ − *ψ*_*A*_ across network ensembles, we need to remove the confounding effect of basin size distribution. We achieve this by assuming a uniform basin size distribution in (0.14):

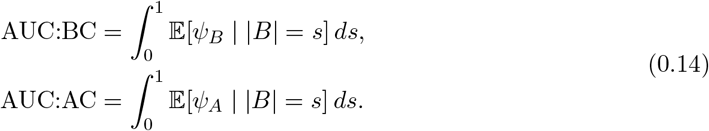

AUC:BC and AUC:AC denote the area under the curves of the functions 𝔼[*ψ*_*B*_ | |*B*| = *b*] and 𝔼[*ψ*_*A*_ | |*B*(*A*)| = *b*], respectively. To compare the basin and the attractor coherence, we consider the *coherence gap*

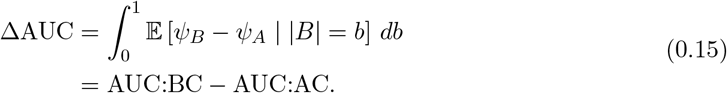

For a network with completely random basin structure, the coherence of an attractor and its corresponding basin are both expected to equal the relative basin size. Therefore, in a truly random network

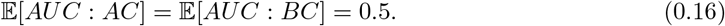

We thus use this value as a baseline for comparison.

In practice, we approximate these integrals on 1000 equal-sized subintervals of [0, 1] with empirical conditional means. Whenever the empirical basin size distribution of a network ensemble exhibits a gap ≥ 0.02, 𝔼[*ψ*_*B*_ | |*B*| = *s*] and 𝔼[*ψ*_*A*_ | |*B*| = *s*] cannot be accurately computed for a non-negligible proportion of relative basin sizes *s*, meaning AUC:BC, AUC:AC and ΔAUC may be incorrect. We therefore did not include these cases in the analysis.

### Bias, sensitivity and dynamical regime

Sensitivity and bias are two key properties that shape the dynamical regime of Boolean networks. The *average sensitivity S*(*f* ) of a Boolean function *f* (*x*,_1_, …, *x*_*n*_) quantifies how sensitive the output is to a change in a single random input [26]. That is,

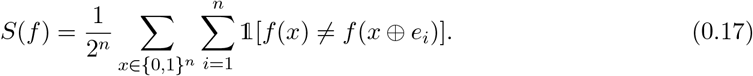

The sum of the average sensitivities of all update rules of a Boolean network is a proxy for its *dynamical regime* [6, 27, *12]*. *Networks with average sensitivity below 1 lie in the ordered dynamical regime* where perturbations vanish over time. Conversely, networks with sensitivity above 1 lie in the *chaotic dynamical regime* where perturbations amplify. Between these two extremes lies the *critical dynamical regime*, associated with an average sensitivity value near 1.

The *bias p* of a Boolean function is defined as the probability of a 1 in the function’s truth table [28]. It quantifies the level of output asymmetry. Functions with *p* = 0.5 are *unbiased*, containing an equal number of zeros and ones. We report bias using two related metrics: *absolute bias a*(*p*) = |2*p* − 1| (ranging from 0 for unbiased to 1 for maximally biased functions) and *standardized bias* ζ(*p*) = *p*(1 − *p*) (ranging from 0 for maximally biased to 0.25 for unbiased functions). The standardized bias and absolute bias are negatively related (Fig. 3A):

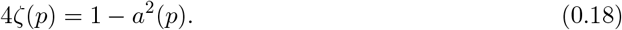

**Figure 3:**
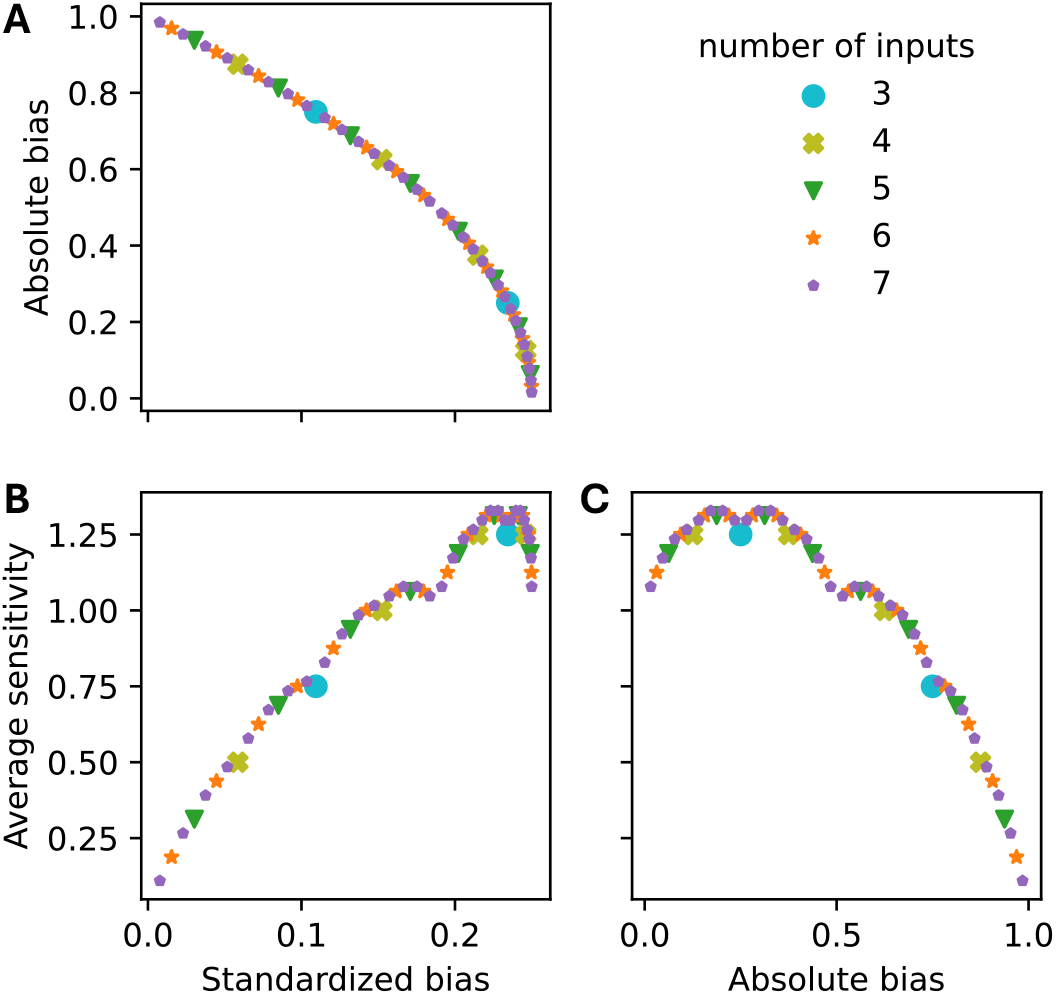
Relationship between standardized bias, absolute bias and average sensitivity for all NCFs with 3-7 inputs. Each dot corresponds to an NCF of defined degree (color) and specific layer structure.

Importantly, bias and sensitivity are not independent: higher bias generally leads to lower sensitivity, though this relationship is non-monotonic [12]. Functions with extreme bias (*p* near 0 or 1) typically exhibit low sensitivity because their outputs are dominated by the preferred value. However, moderately biased functions exhibit varying sensitivity, particularly depending on their degree of canalization.

### Canalization

Canalization was introduced in developmental biology to describe the regulative behavior of developmental pathways, i.e., their tendency to return to a normal trajectory despite alterations in their state [1]. Consider a Boolean function *f* (*x*_1_, …, *x*_*n*_) of degree *n*. The variable *x*_*i*_, 1 ≤ *i* ≤ *n*, is a *canalizing variable* if there exists a fixed value *a* ∈ {0, 1} such that when *x*_*i*_ = *a*, the output of *f* is fully determined, independently of the other inputs [29]. The function *f* is *canalizing* if it has a canalizing variable. This definition formalizes the idea of canalization by describing how specific variables can enforce stability in developmental pathways by effectively diminishing the impact of other influences.

Canalizing Boolean functions can be further classified by their *(canalizing) layer structure*, which can be computed iteratively [30]. First, identify all *k*_1_ canalizing variables and their corresponding canalizing values. These variables and their values form the first canalizing layer. To determine the next layer, substitute the non-canalizing values into the function. This results in a new function that depends only on the variables not included in the first layer. Then, identify the new function’s canalizing variables and their corresponding canalizing values. These form the second layer and their number is denoted by *k*_2_. Repeat this process until either a constant function or a non-canalizing function is reached after *r* ≥ 0 steps. The vector (*k*_1_, …, *k*_*r*_) constitutes the canalizing layer structure of the original function *f* [12, 31]. The *canalizing depth* of *f* is 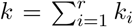 and describes the number of variables that become canalizing eventually. Note that, a Boolean function is canalizing when the canalizing depth *k* ≥ 1. *Nested canalizing functions* (NCFs) are Boolean functions whose canalizing depth equals their degree. In other words, all the inputs of NCFs become eventually canalizing. Most updates rules in biological BNs are NCFs [10].

To illustrate the concept of layer structure, consider the 3-input Boolean functions *f* (*x, y, z*) = *x*∧*y* ∧*z* and *g*(*x, y, z*) = *x*∧ (*y* ∨*z*). Both functions are nested canalizing, but they exhibit different dynamical behavior. The function *g* is more sensitive than *f*, and this difference arises from their distinct layer structures. In *f*, all three variables are canalizing. Setting any one of them to zero immediately determines the output of the function. Therefore, all variables belong to a single canalizing layer, and the layer structure of *f* is (3). In contrast, only *x* is canalizing in *g*. When *x* is set to its non-canalizing value, the function reduces to *y* ∨*z*, where both *y* and *z* become canalizing. Thus, the layer structure of *g* is (1, 2).

NCFs with the same layer structure have the same average sensitivity and absolute bias [12]. For example, the function *f* has absolute bias 3*/*4 because its truth table contains either seven ones. Similarly, the function *g* contains five ones and thus has absolute bias 1*/*4. For a given number of variables *n* ≥ 2, there are 2^*n*−2^ NCFs with different layer structure, each appearing equally likely by chance. Table 1 illustrates this for *n* = 5, showing how layer structure determines both bias and sensitivity – a relationship we exploit in our results.

**Table 1:** Stratification of all 5-input NCFs by layer structure, similar to [12].

| Layer structure | Example | Hamming weight | Absolute bias | Standardized bias | Average sensitivity |
| --- | --- | --- | --- | --- | --- |
| (5) | $A \vee B \vee C \vee D \vee E$ | 1 or 31 | 15/16 | 0.0303 | 0.3125 |
| (3, 2) | $A \vee B \vee C \vee (D \wedge E)$ | 3 or 29 | 13/16 | 0.0850 | 0.6875 |
| (2, 1, 2) | $A \vee B \vee (C \wedge (D \vee E))$ | 5 or 27 | 11/16 | 0.1318 | 0.9375 |
| (2, 3) | $A \vee B \vee (C \wedge D \wedge E)$ | 7 or 25 | 9/16 | 0.1709 | 1.0625 |
| (1, 1, 3) | $A \vee (B \wedge (C \vee D \vee E))$ | 9 or 23 | 7/16 | 0.2021 | 1.1875 |
| (1, 1, 1, 2) | $A \vee (B \wedge (C \vee (D \wedge E)))$ | 11 or 21 | 5/16 | 0.2256 | 1.3125 |
| (1, 2, 2) | $A \vee (B \wedge C \wedge (D \vee E))$ | 13 or 19 | 3/16 | 0.2412 | 1.3125 |
| (1, 4) | $A \vee (B \wedge C \wedge D \wedge E)$ | 15 or 17 | 1/16 | 0.2490 | 1.1875 |

For a broader treatment and alternative definitions of canalization in BNs, we refer the reader to [32, 33, 34, 35]. In these papers, the traditional definition of canalization by Kauffman is generalized to the *canalizing strength*, which quantifies the level of collective canalization in a Boolean function [34]. Canalization can also be interpreted as *input redundancy*, which is the probability that an input to a Boolean function is not needed to determine its output [33, 35]. By contrast, the *effective degree* describes the average number of inputs needed to determine the output.

In this paper, we present evidence that the gap between attractor and basin coherence widens as the level of canalization in the network increases, irrespective of how canalization is assessed.

### Analysis of biological network models

In this study, we considered two collections of BN models: synthetic random networks and published biological models. The latter is a collection of 122 distinct models that contain between 5 and 342 nodes [10]. For small BNs, the entire state transition graph, basin coherences, and basin sizes can be computed quickly. However, as the number of nodes increases, the exponential growth of the state space makes it infeasible to compute basin sizes and basin coherences exactly. Therefore, we conducted two types of biological model analyses: one for smaller networks (*N* ≤ 20) and another for larger networks (*N* ≤ 64).

For the first type of analysis, we considered the 42 out of the 122 biological BN models with *N* ≤ 20 nodes. For these networks, we computed the entire state transition graph for up to 16 different fixed-source networks as described below. Consequently, we obtained exact values for the basin sizes, basin coherences *ψ*_*B*_, and attractor coherences *ψ*_*A*_.

For the second type of analysis, we considered 95 models with *N* ≤ 64. For these networks, we approximated the state transition graph using BoolForge [36]. We generated *q* = 1000 random initial states **x** ∈ {0, 1}^*N*^ and repeatedly updated **x** until it transitioned into a periodic orbit, i.e., to an attractor. The proportion of random initial states that transition to a specific attractor provides an unbiased estimate for the corresponding relative basin size. Similarly, the coherence of these states offers an unbiased estimate of basin coherence. This procedure identifies an attractor of relative basin size *b* ∈ (0, 1] with probability *p*(*b, N* ) = 1−(1−*b*)^*q*^ [37]. Consequently, attractors with very small basins are often overlooked. However, biologically meaningful attractors typically have sufficiently large basin sizes and are discovered with high probability. For example, an attractor with relative basin size *b* = 1% is found with probability 99.996%. Irrespective of the network size, we computed the exact attractor coherence *ψ*_*A*_ for every identified attractor *A*. Thus, for this analysis, we obtained the exact value of *ψ*_*A*_ and unbiased estimates of *ψ*_*B*_ and basin sizes for all the identified attractors.

### Treatment of source nodes

Some of the biological network models contain *source nodes* – variables with no regulatory inputs whose values remain constant throughout the dynamics [16]. For a network with *M* source nodes, the full state transition graph of size 2^*N*^ decomposes into 2^*M*^ independent subgraphs, each corresponding to one fixed configuration of source node states. Each subgraph evolves under identical regulatory logic for the remaining *N* − *M* nodes but with different constant inputs from the sources. This partitioning can artificially inflate the number of attractors and deflate basin sizes, confounding comparisons of coherence between networks that differ in the number of source nodes. To remove this effect, we analyzed only *fixed-source networks* in the biological datasets, where the states of the source nodes were held constant during the dynamical simulations. For networks with multiple source nodes, we randomly analyzed up to 16 of the 2^*M*^ independent state transition graphs and average the resulting attractor statistics. This approach isolates the contribution of the regulatory logic from trivial fragmentation effects introduced by variable numbers of source nodes.

### Analysis of random Boolean networks

Random models offer a useful baseline for interpreting how various features of biological networks shape their dynamics. We conducted two experiments on random ensembles of BNs to examine how canalization influences the relationship between attractor and basin coherence. In both experiments, each ensemble consisted of 10,000 randomly generated 12-node networks, created and exactly analyzed using the Python package BoolForge [36]. In the first experiment, all networks within an ensemble share update rules with the same degree and canalizing depth, while these parameters vary across different ensembles. We ensured that (i) the wiring diagrams of all networks were strongly connected and (ii) the Boolean functions were non-degenerate (i.e., function outputs genuinely depend on all input variables indicated in the wiring diagram, not a subset). This excludes, for example, *f* (*x, y, z*) = *x* ∨ *y* as a 3-input update function. In the second experiment, all networks within an ensemble were governed by the same type of NCF, specified by canalizing layer structure.

## Results

### Basin and attractor coherence in biological networks

A recent meta-analysis of 122 published, expert-curated biological BN models revealed that they possess a number of remarkable structural and dynamical features [10]. They are sparsely connected and the Boolean rules describing the regulatory logic are significantly more canalizing, redundant and biased than expected by chance. To assess how these features specifically influence the robustness of biological BNs, we first considered the 42 models of size *N* ≤ 20. These models are small enough to be simulated exhaustively, allowing us to compute their coherence exactly.

For each of the 42 biological models, we generated 100 corresponding random null models. These null models preserve the wiring diagram but used randomized, non-degenerate update rules. To examine the difference in coherence between the null models and the biological models, we computed the difference between the coherence of the biological networks and the mean coherence of the corresponding random models. The distribution of paired coherence differences did not deviate from normality (Shapiro–Wilk *p* = 0.31), and a one-sided paired t-test confirmed that biological networks exhibited significantly higher coherence than the random null models with the same topology (*p* = 9 *×* 10^−6^, *n* = 42; Fig. 4). This demonstrates that the high coherence in biological networks is not solely due to their network topology but likely arises from their specific regulatory logic.

**Figure 4:**
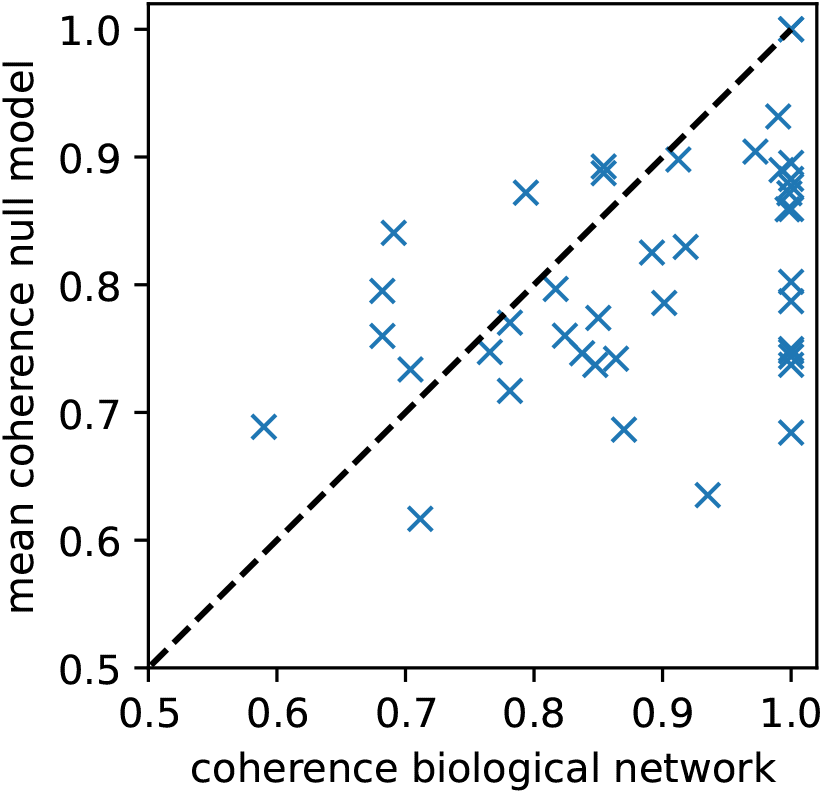
Biological Boolean network models exhibit higher-than-expected coherence. For each of the 42 biological BN models from [10] with size *N* ≤ 20, the network coherence is plotted against the mean coherence of 100 corresponding random null models. The null models have the same wiring diagram but use randomized, non-degenerate update rules. The dashed diagonal line indicates equality between biological and expected coherence.

To further investigate the mechanisms underlying this enhanced stability, we examined the coherence of basins and attractors in biological networks. For each network attractor, we computed the basin size as well as the basin and attractor coherence. Across all attractors, larger basins are generally more robust (Fig. 5), and basin and attractor coherence are, not surprisingly, highly correlated (Pearson’s r = 0.97). It is known that BNs with low connectivity have high average basin coherence, while highly connected BNs possess basins that are almost randomly distributed throughout the state space [14]. Given the relatively low connectivity of the biological BN models (mean=2.56, median=2.27) [10], this helps explain why even attractors with relatively small basins were on average fairly coherent.

**Figure 5:**
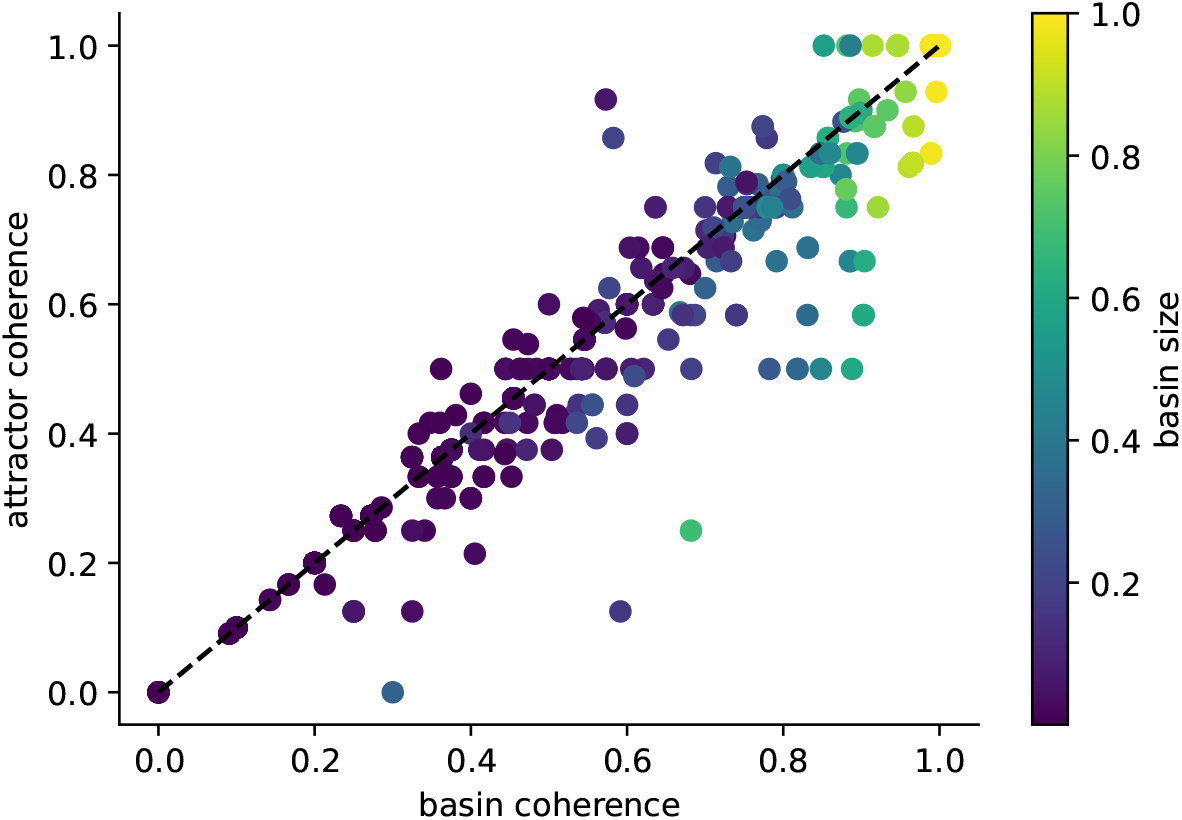
Exact basin and attractor coherence of biological Boolean network models. For each attractor found in Boolean biological network models in [10] of size *N* ≤ 20 (42 models), attractor coherence is plotted against basin coherence. Color indicates the relative basin size. A substantial number of attractors lie below the diagonal (basin coherence *>* attractor coherence; sign test *p* = 2 *×* 10^−11^), indicating that attractors are on average less resilient to perturbations than their basins.

We observed a substantial number of attractors with high basin coherence but relatively low attractor coherence (sign test, used because the distribution of differences is non-normal and asymmetric: *p* = 2 *×* 10^−11^, *n* = 597; Fig. 5). This pattern suggests that some attractors lie near the “boundary” of their basin. It further implies that the system typically remains within the same basin following perturbations of transient states, whereas perturbations applied at an attractor more frequently redirect the system to a different attractor. This pattern partially persists when sampling the dynamics of 95 biological BN models of size *N* ≤ 64 (Fig. 6). For attractors with relatively large, biologically relevant basins, the average basin coherence is higher than the average attractor coherence. In contrast, attractors with tiny basins exhibit on average the opposite trend. Altogether, the area under the basin coherence curve (AUC:BC = 0.853) is larger than the area under the attractor coherence curve (AUC:AC = 0.825). This is consistent with the landscape model illustrated in Fig. 1, where attractors occupy positions near relatively flatter boundaries of their basins. To disentangle what features of biological networks render their attractors less robust than their basins, we next investigated large ensembles of random BNs with a fixed in-degree.

**Figure 6:**
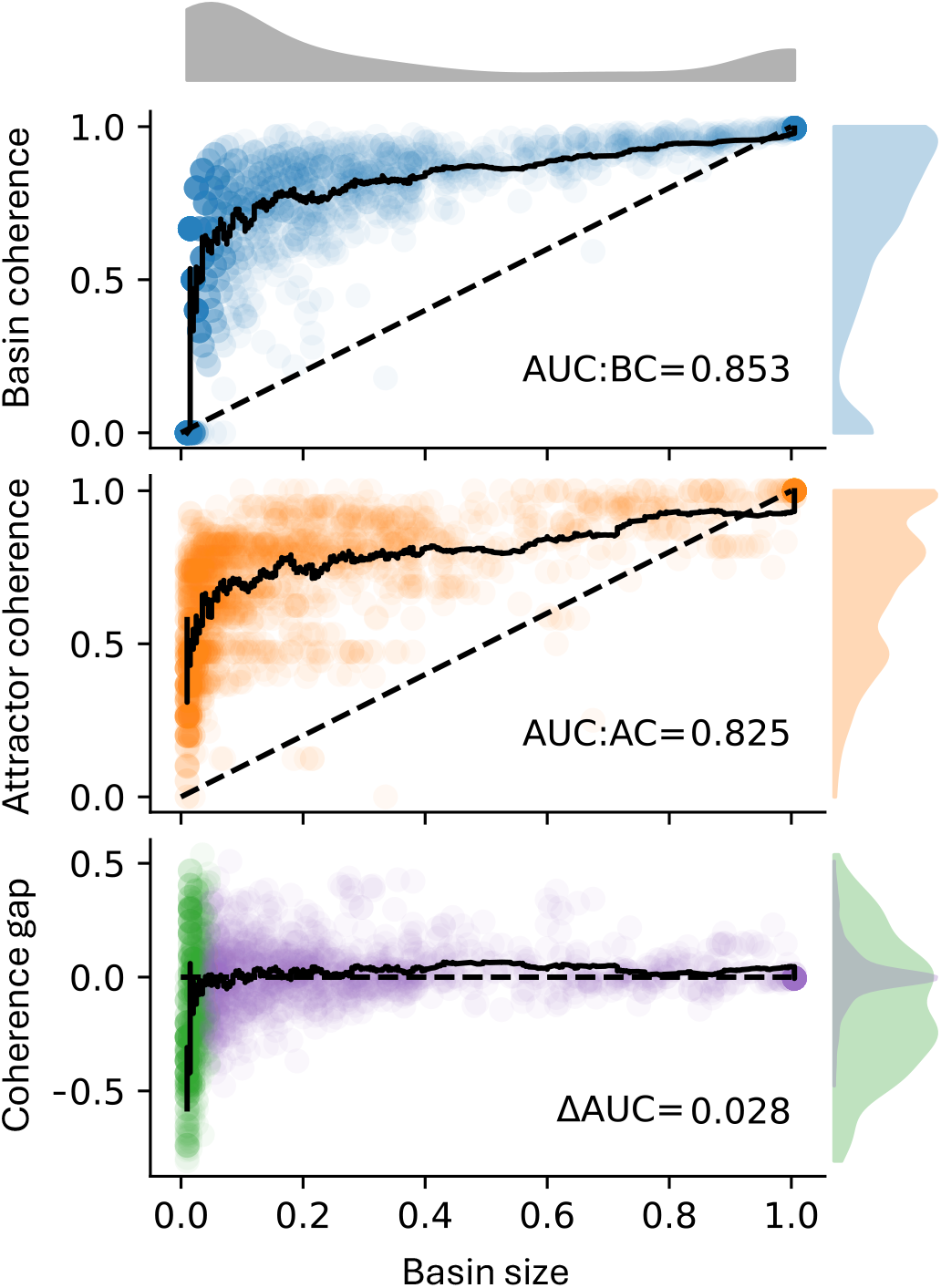
Coherence of biological Boolean network attractors and their basins. The synchronous state transition graph (attractors, basin coherence, attractor coherence) was approximated for all Boolean biological network models in [10] of size *N* ≤ 64 (95 models, see Methods for details). The approximated basin coherence (top; blue), the exact attractor coherence (middle; orange) and the difference per attractor (bottom) are plotted against the approximate basin size (x-axis) for each discovered attractor in the biological networks. Black lines depict an unweighted rollingwindow mean with window size 50. Distributions of the different metrics are color-coded. The distribution of the difference in basin minus attractor coherence is shown for basin sizes *<* 5% (green) and ≥ 5% (purple). AUC quantifies the area under the respective lines.

### Canalization increases both basin and attractor coherence but creates coherence gap

One key feature of biological BN models is their high level of canalization [18, 8, 35, 10]. To assess the effect of canalization on the basin and attractor coherence, we generated ensembles of 10,000 12-node BNs for different combinations of degree and canalizing depth (Fig. 7 and Fig. 8).

**Figure 7:**
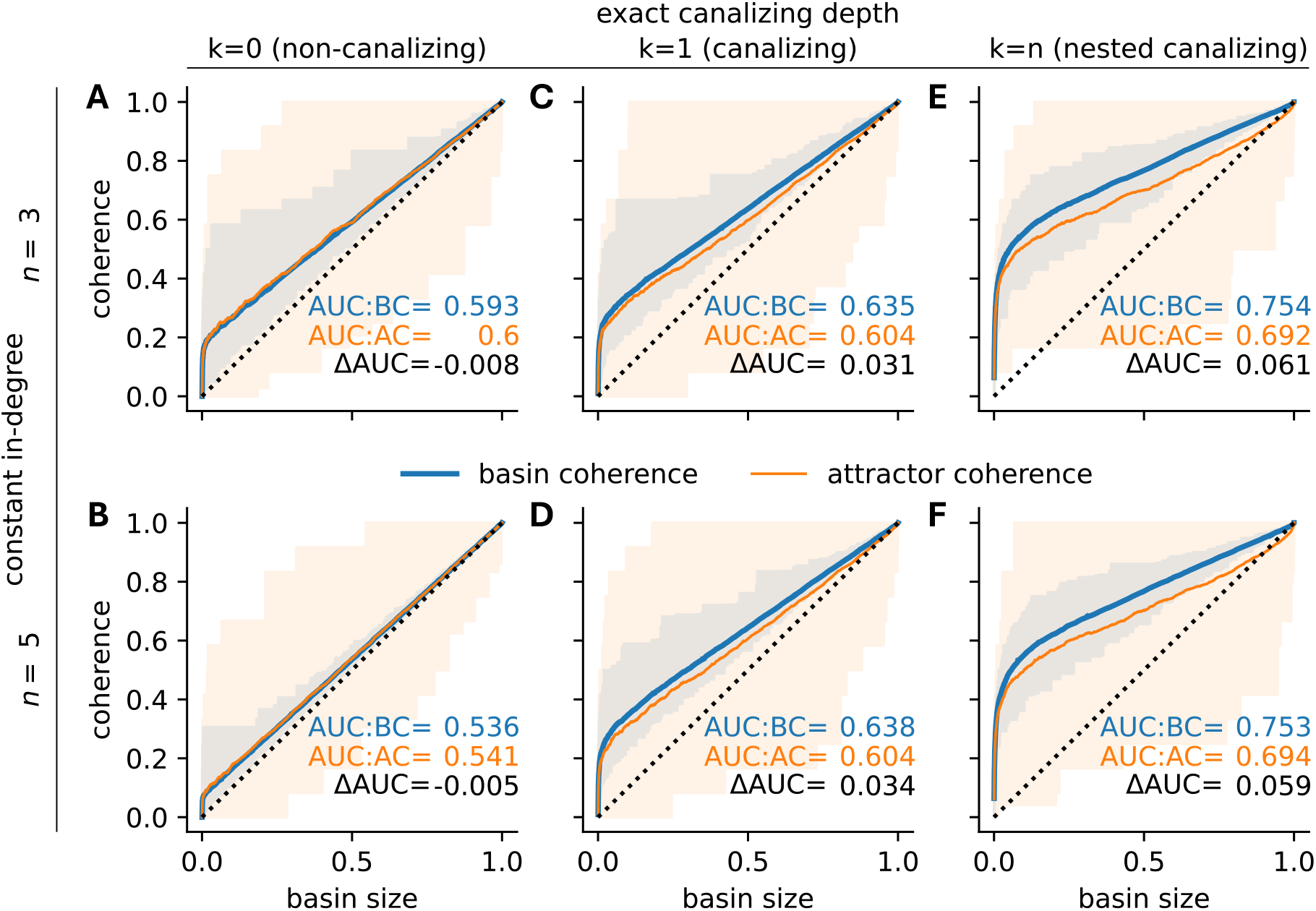
Robustness of random Boolean network attractors and their basins. For 10,000 random 12-node BNs with random (A,B), canalizing (C,D) and nested canalizing (E,F) update rules of in-degree 3 (A,C,E) and 5 (B,D,F), the coherence of each attractor (orange) and the coherence of its corresponding basin (blue) are stratified by basin size (x-axis). Shaded areas indicate the range of observed values. Lines depict rolling-window means with window size 500 and AUC quantifies the area under the respective lines.

**Figure 8:**
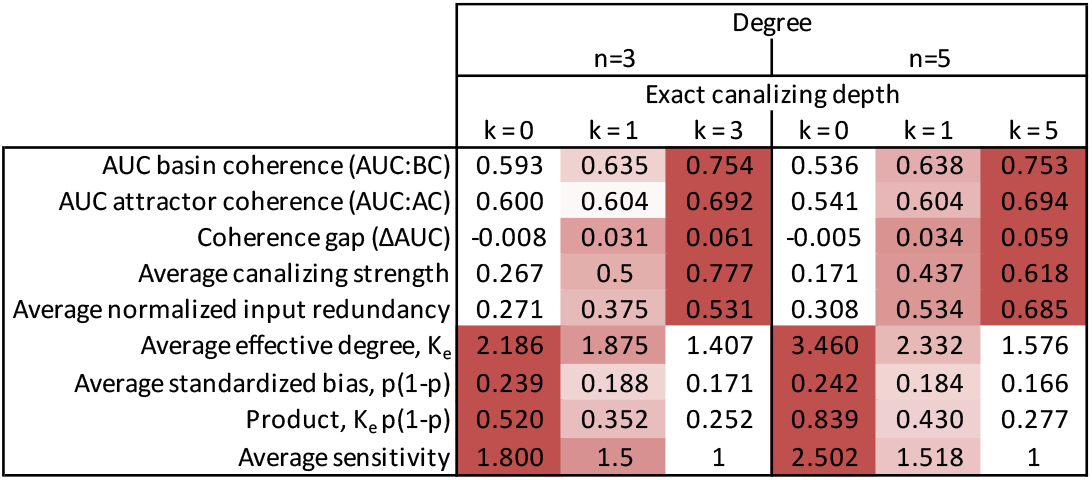
Properties of non-canalizing, canalizing and nested canalizing Boolean functions and networks governed by them. For each metric and a given degree, a separate linear color gradient is used.

In the absence of canalization, increasing the degree led to randomly distributed basins in the state space. That is, both basin and attractor coherence converged toward the random baseline given by the relative basin size. For example at degree 3, AUC:BC and AUC:AC values were 18.6% and 20% above the random baseline of 0.5 (Fig. 7A). These values dropped to 7.2% and 8.2% at degree 5 (Fig. 7B). In contrast, canalizing and nested canalizing networks exhibited substantially higher basin and attractor coherence, regardless of degree. Specifically, the AUC:BC values were 27% and 27.6% higher in canalizing networks of degree 3 and 5 (Fig. 7C,D). In nested canalizing networks, these numbers rose to 50.8% and 50.6%, respectively (Fig. 7E,F). These observations are consistent with earlier findings that biological networks, typically highly canalizing, tend to be highly coherent.

Canalization increases both basin and attractor coherence in absolute terms, but increases basin coherence more strongly, creating a coherence gap. This implies that transient states in canalizing and nested canalizing networks are generally more coherent than attractor states, while there exists hardly any difference in non-canalizing networks. For canalizing networks, AUC:AC values were only 20.8% higher than the random baseline (Fig. 7C,D), representing a relative drop (ΔAUC*/*(AUC:BC − 0.5)) of 22.9% (n=3) and 25.2% (n=5), compared to the AUC:BC values. In nested canalizing networks, AUC:AC values were 38.4 to 38.8% higher than the baseline, corresponding to a similar relative drop of 24.4% (n=3) and 23.3% (n=5). Biological BNs are often dominated by NCFs [10], which may explain the observed coherence gap in biological systems.

There are several alternative measures of canalization in Boolean functions, including canalizing strength and input redundancy [35, 34]. In this study, we chose canalizing depth as our primary metric because the standard monomial form of a Boolean function provides a straightforward method for generating networks with a specified canalizing depth [30]. Functions with higher canalizing depth tend to exhibit higher canalizing strength and input redundancy (Fig. 8). These high correlations suggest that the findings are insensitive to the specific choice of canalization metric.

### Sensitivity drives basin coherence

Sensitivity and bias are two key network properties that shape a network’s dynamical behavior. Both properties are closely linked to canalization. In particular, canalizing functions tend to be biased [19], and they typically exhibit low sensitivity. Furthermore, other well-established predictors of a network’s dynamical regime, such as the metric ⟨*K*_*e*_⟩⟨*p*(1−*p*)⟩, which combines average effective degree and average standardized bias, are also highly correlated with canalization [38, 10]. These relationships suggest that the dynamical regime of a network, shaped by canalization and bias, may play a key role in determining the coherence of its basins and attractors (Fig. 8).

To systematically dissect the roles of sensitivity and bias in shaping the observed coherence gap, we re-analyzed the networks governed by NCFs (Fig. 7E,F). To achieve finer resolution, we further stratified NCFs according to their canalizing layer structure. Because NCFs with the same layer structure exhibit identical bias and average sensitivity [12], this approach allowed us to independently examine how sensitivity and bias influence basin coherence, attractor coherence, and the coherence gap. Specifically, for each 5-input NCF with different layer structure, we generated an ensemble of 10,000 12-node BNs, yielding networks with varying levels of sensitivity and bias (Table 1).

Basin coherence was high across all network ensembles, but showed clear sensitivity dependence (Fig. 9). Networks governed by NCFs with the highest average sensitivity of 1.3125 – those with layer structures (1, 1, 1, 2) and (1, 2, 2) – exhibited the lowest AUC:BC values (Table 1, Fig. 9F,G). In contrast, bias proved to be a weak predictor of basin coherence. For example, NCFs with layer structure (1,4), which are nearly unbiased (absolute bias = 1/16), and NCFs with layer structure (3,2), which are highly biased (absolute bias = 13/16), both exhibited similarly high AUC:BC values. These findings suggest that the sensitivity plays a more important role than bias in determining AUC:BC values.

**Figure 9:**
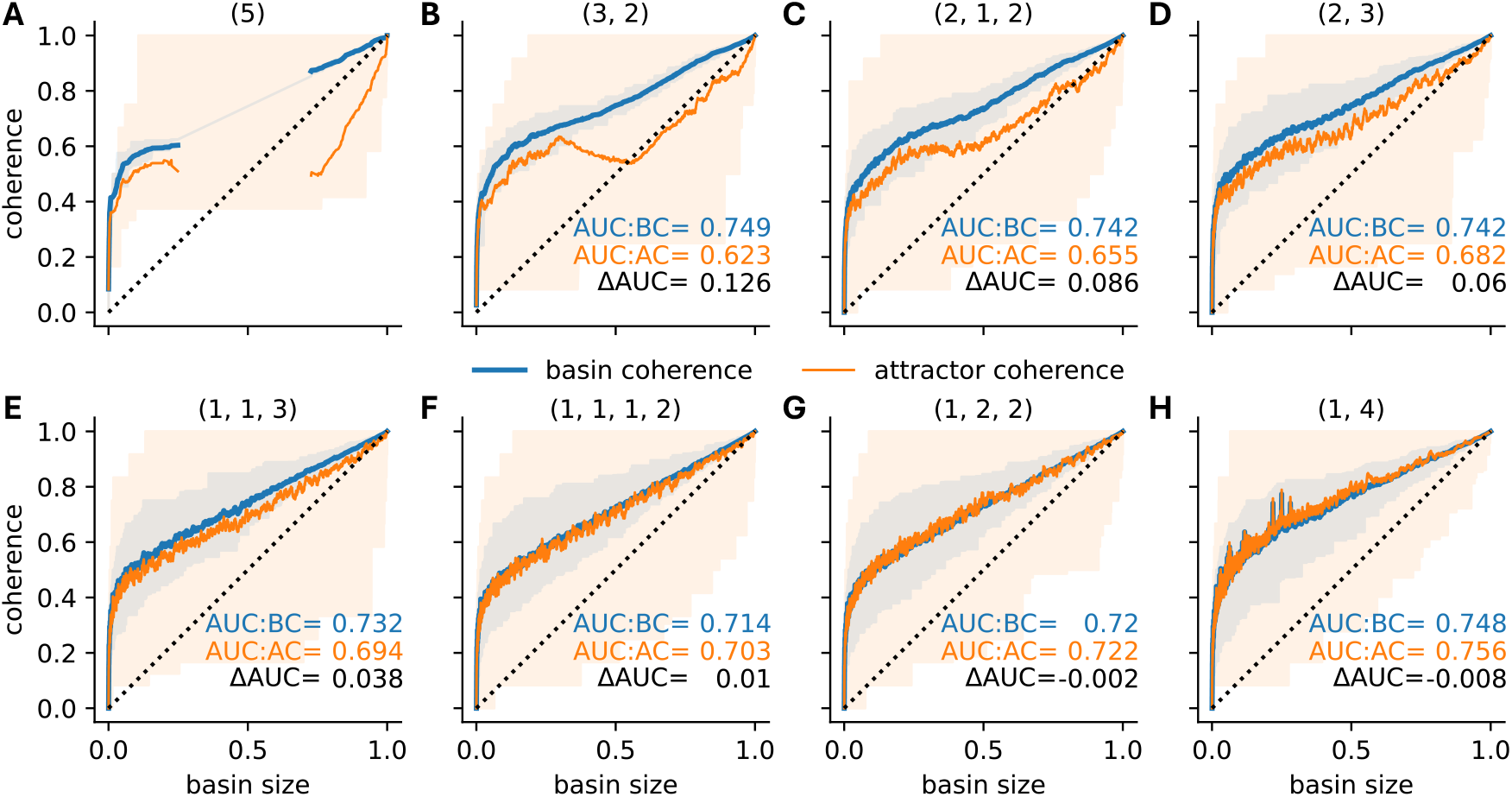
Robustness of nested canalizing Boolean network attractors and their basins. For 10,000 random 12-node BNs governed by 5-input NCFs of different layer structure (specified above each panel), the coherence of each attractor (orange) and the coherence of its corresponding basin (blue) are stratified by basin size (x-axis). Shaded areas indicate the range of observed values. Lines depict rolling-window means with window size 50 and AUC is the area under the respective lines. The basin size distribution corresponding to each type of network is shown in Fig. S1.

Repeating this analysis for networks ensembles with different connectivity (3-7 inputs per node) revealed a consistent pattern. AUC:BC values were consistently lowest in networks governed by the most sensitive NCFs (Fig. 10A). However, the observed relationship between AUC:BC and the average sensitivity was far from monotonic (Spearman’s *ρ* = -0.629): Critical networks (sensitivity ≈ 1) exhibited locally minimal basin coherence, while the most ordered networks (lowest sensitivity) did not possess maximal AUC:BC values. Similar trends persisted when comparing AUC:BC values with the standardized bias of the NCFs (Fig. 10B; Spearman’s *ρ* = -0.312), with one notable exception – networks governed by the least biased NCFs maintained relatively high basin coherence. This exception is in accordance with the underlying non-monotonic relationship between bias and sensitivity (Fig. 3B,C): NCFs with low absolute bias exhibited lower sensitivity values.

**Figure 10:**
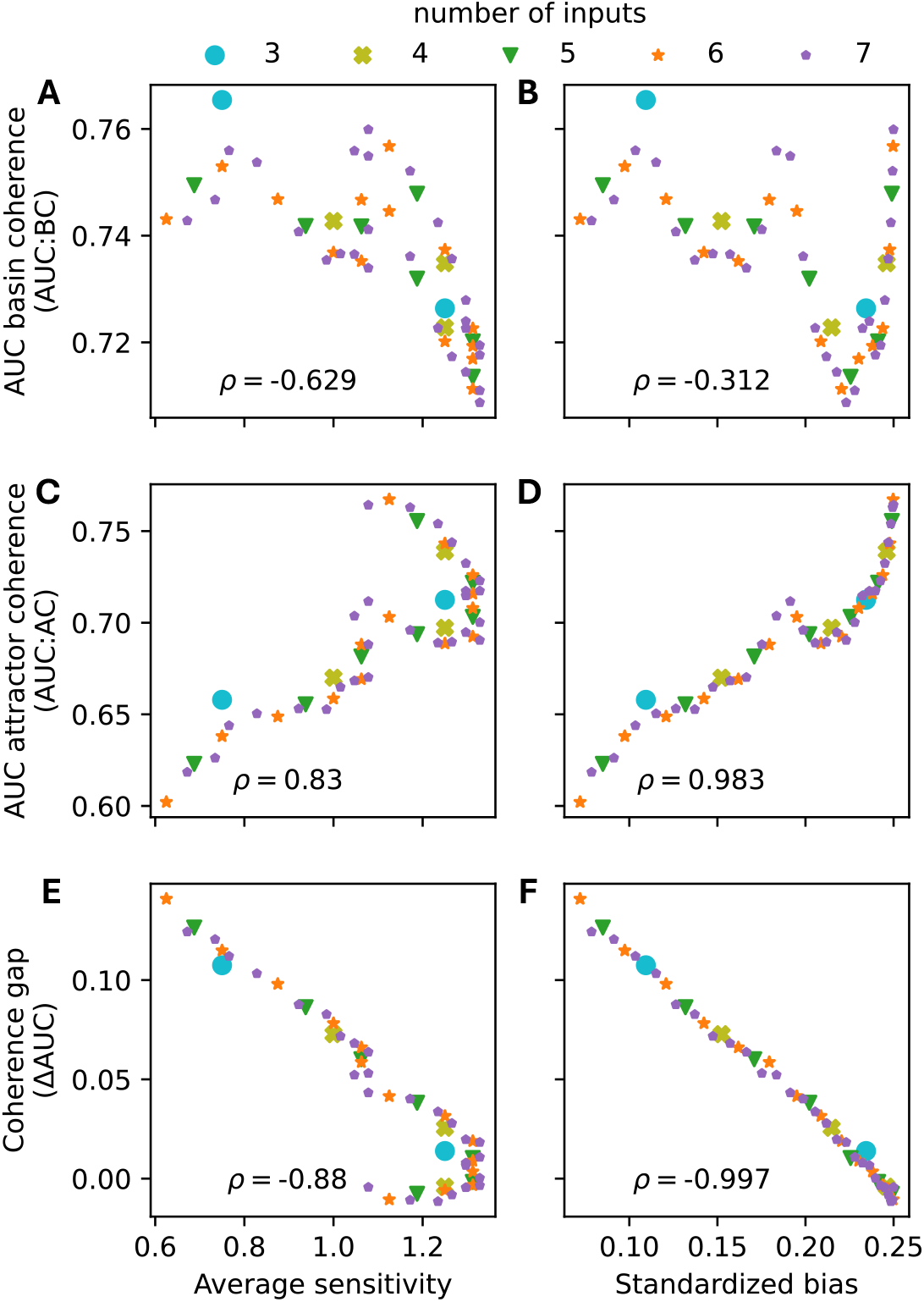
Association between different metrics. For ensembles of random 12-node BNs governed by nested canalizing update rules in 3-7 inputs (color) and of different layer structure, AUC:BC (top row), AUC:AC (middle row) and the coherence gap (bottom row) are computed as in Fig. 9, and plotted against the average sensitivity (left column) and the standardized bias *p*(1 − *p*) (right column). Each data point is derived from 10,000 networks. Across all data points, Spearman’s *ρ* correlation coefficient is computed.

### Bias drives attractor coherence

In contrast to basin coherence, attractor coherence varied substantially across network ensembles and depended more strongly on bias than sensitivity (Fig. 10C,D). Generally, the lower the standardized bias of the NCFs (i.e., the higher the absolute bias), the lower was the observed attractor coherence (Spearman’s *ρ* = 0.983). This suggests that, in networks governed by more biased NCFs, attractors with small and moderately-sized basins are less coherent. Such attractors are, however, also less frequent in these biased networks (Fig. S1, Figs. **??**,**??**).

Intriguingly, the attractor coherence of networks in the ordered dynamical regime, governed by NCFs with layer structure (3, 2) and (2, 1, 2) with average sensitivity below 1 (i.e., with ordered dynamics), appeared to be non-monotonic in basin size (Fig. 9B,C). For example, attractors in networks governed by NCFs with layer structure (3, 2) that attract roughly half the states were on average less coherent than attractors with a basin size of around 30%. More connected networks governed by 6-input and 7-input NCFs exhibited the same trends, indicating robust results (Figs. S2, S3, S4, S5).

### Bias perfectly explains the coherence gap

The coherence gap exhibited a remarkably strong linear relationship with standardized bias (Spearman’s *ρ* = -0.997;Fig. 10F). Networks governed by least biased NCFs (with standardized bias *p* ≈ 0.25) possessed roughly equal attractor and basin coherence. As the standardized bias decreases (i.e., as the absolute bias increases), the coherence gap increased nearly linearly. This pattern held across all investigated in-degrees and suggests that bias, modulated through canalization [19, 12], largely determines whether, and by how much, attractor and basin coherence differ. We hypothesize that this arises because highly biased networks confine attractors to similar regions of state space. In these networks, most nodes become frozen in the same state across attractors, with dynamics confined to a small non-frozen core [39]. This clustering reduces the mean Hamming distance between attractors relative to random state pairs, creating relatively flatter basin boundaries near attractors compared to mid-trajectory states, while maintaining high absolute coherence throughout the basin. This provides the mechanism underlying the within-basin stability gradient shown in Fig. 1: bias-driven compression of attractors into similar state-space regions creates the coherence gap. The biological relevance of this gradient depends on whether multiple attractors with substantial basins coexist (Fig. S1).

## Discussion

Our analyses demonstrate that biological Boolean networks exhibit markedly higher coherence than random ensembles of comparable size and connectivity. This enhanced resilience to perturbations can be traced back to characteristic design principles of biological systems, such as abundant canalization and high bias. Each of these properties has been observed consistently across curated biological network models and contributes to systematic dynamical differences between biological and synthetic networks [27, 8, 40, 10].

### Attractors are less stable than their basins

A central result of this study is the discovery that attractor coherence is lower than basin coherence in biological networks. This finding substantially revises our understanding of phenotypic stability. Rather than occupying the most stable positions in their regulatory landscapes, mature cell types and phenotypes are positioned closer to basin boundaries relative to transient states, despite maintaining high absolute coherence. This within-basin stability gradient means that perturbations of attractor states are more likely to induce phenotype switching than perturbations of transient states in the same basin.

### Canalization creates the coherence gap through bias

Our systematic analysis of random network ensembles reveals the mechanism underlying this phenomenon. Canalization – a pervasive feature of biological regulation where specific inputs can override others – simultaneously increases overall network stability and magnifies the coherence gap. This dual effect arises because canalization induces strong bias in Boolean functions [19, 12]. The coherence gap is almost perfectly predicted by network bias (Spearman’s *ρ* = −0.997), with highly biased networks showing the largest disparities between basin and attractor coherence. One possible explanation is that attractors in highly biased and canalized networks typically cluster in similar regions of the state space. This occurs because bias drives many nodes to become frozen (always 0 or always 1), leaving dynamics confined to a small non-frozen core [39]. With attractors concentrated in similar regions, they sit closer together in state space, reducing the mean Hamming distance between attractors and effectively flattening the ridges separating basins near their valley floors. Small perturbations at these attractor locations can thus more easily induce inter-attractor transitions.

Importantly, this relationship holds across multiple measures of canalization – including canalizing depth [30], canalizing strength [34], and input redundancy [35] – indicating robustness of our findings to methodological choices. The dynamical regime (ordered, critical, or chaotic) plays a secondary role, primarily influencing basin coherence while bias dominates attractor coherence.

### Context-dependent relevance of the coherence gap

While canalization increases both basin and attractor coherence in absolute terms – consistent with the high robustness of biological systems – the coherence gap is a relative phenomenon describing the within-basin stability gradient. Its biological relevance depends strongly on the basin size distribution.

In highly biased, canalized networks, the number of attractors is often small (Fig. S1). Networks may be dominated by one or a few major attractors occupying most of state space. In such cases, the coherence gap has limited functional impact because alternative phenotypes have been effectively or completely eliminated.

However, the coherence gap becomes functionally important in biological contexts where multiple phenotypically meaningful attractors coexist. In these contexts, our finding that attractors are less stable relative to their basins predicts enhanced phenotypic plasticity despite overall high stability. The impact of canalization is thus context-dependent: protective when one fate dominates (high absolute coherence, coherence gap irrelevant), permissive when choices exist (high absolute coherence, but the coherence gap facilitates phenotypic transitions). This dual nature may help resolve the apparent paradox of how biological systems can be simultaneously robust and adaptable.

### Balancing robustness and adaptability: insights into the criticality hypothesis

These findings provide new insight into the longstanding robustness-adaptability trade-off in biological systems [7, 41, 9]. Park et al. (2023) demonstrated that biological networks are substantially more robust than previously appreciated when source nodes are properly accounted for, challenging the criticality hypothesis [16]. Our results refine and extend this picture: Since biological networks rest at attractors, their attractor coherence primarily determines their resilience to perturbations. Since biological networks possess lower attractor than basin coherence indicates, they operate in fact closer to the critical edge.

Canalization and bias create a dual-purpose architecture operating at two levels. First, at the level of absolute stability, both basin and attractor coherence are high, ensuring that random perturbations rarely cause phenotype switching. This explains developmental robustness – stochastic fluctuations do not derail differentiation, and mature cell types resist random noise. Second, within each basin, a stability gradient positions attractors closer to boundaries relative to transient states. This relative instability is dormant when only one attractor dominates but becomes functionally relevant when multiple fates are accessible, enabling regulated phenotype transitions in response to appropriate signals.

This architecture may be evolutionarily advantageous. During normal development, high absolute coherence protects cells from stochastic perturbations as they follow canalized paths through state space. The basin’s depth ensures reliable lineage commitment. However, when cells reach decision points where multiple fates have substantial basins, the coherence gap provides a mechanism for controlled plasticity: Cells can respond to developmental signals, injury, or stress by transitioning to alternative fates without requiring dramatic rewiring of regulatory networks. The system is robust by default but plastic on demand.

### Implications for development, disease, reprogramming, and design

The attractor-centric perspective introduced here has broad implications across biological and engineering contexts.

### Development and cell fate decisions

High basin coherence during differentiation ensures robust lineage commitment, while the within-basin stability gradient means attractors are less stable than transient states. At developmental branch points where multiple fates remain accessible, the coherence gap predicts that mature differentiated cells should be more susceptible to fate-switching perturbations than cells mid-differentiation. This generates testable predictions: time-resolved reprogramming experiments should show that mature cells respond more readily to reprogramming factors than cells caught mid-differentiation, and developmental stages where multiple fates have comparable basin sizes should exhibit higher phenotypic plasticity.

### Cancer and disease

In normal tissues, regulatory networks typically produce one dominant cellular phenotype. However, in cancer, mutations can create multiple partially stable phenotypic states – epithelial, mesenchymal, and hybrid intermediates – each with non-negligible basin sizes [2, 21, 22]. When such multi-attractor landscapes emerge, our results predict enhanced phenotypic plasticity: cancer cells more readily undergo epithelial-mesenchymal transition, dedifferentiation, or therapyinduced state switching.

### Cellular reprogramming

The relative fragility of attractors provides a mechanistic explanation for why reprogramming of mature cells is possible. Differentiated cells, although occupying “deep valleys” with high absolute coherence, often sit near basin boundaries that can be breached by suitable perturbations, such as transcription-factor cocktails [42, 43]. This framework makes counterintuitive predictions: perturbations applied to fully differentiated cells may induce reprogramming more efficiently than perturbations applied during differentiation, and highly canalized cell types may, paradoxically, be easier to reprogram from their mature state than from intermediate differentiation stages.

### Synthetic biology and bioengineering

For the design of synthetic gene circuits, our results highlight a trade-off between robustness and controllability: canalization increases absolute coherence but creates coherence gaps when multiple functional states exist. For single-state circuits (biosensors, memory devices), high canalization creates a dominant attractor with high stability. For multistate circuits (toggle switches, oscillators), moderate canalization provides intermediate stability for all states, while high canalization creates high absolute stability but positions each state closer to its basin boundary. This suggests maximizing canalization for single-state circuits and tuning it moderately for multi-state circuits requiring balanced robustness across states.

### Limitations and future directions

The coherence gap metric (ΔAUC) removes confounding effects of basin size distribution but may overestimate the importance of the coherence gap in highly ordered networks where mid-sized basins (which exhibit the highest coherence gap) are rare. Alternative metrics that weight by biological relevance or phenotypic importance could provide complementary perspectives.

Additionally, synchronously updated Boolean networks, while capturing key regulatory logic, necessarily abstract away quantitative details of gene expression dynamics, post-translational modifications, and metabolic constraints. Extensions to asynchronously updated, continuous or multi-state models could reveal additional nuances of the stability gradient within basins.

Moreover, developing a deeper theoretical understanding of how bias geometrically constrains attractor positions in state space remains an important direction. While our empirical results demonstrate the near-perfect predictive power of bias, a rigorous mathematical proof connecting bias to attractor boundary proximity would strengthen the mechanistic foundation.

### A revised landscape metaphor

Waddington’s landscape has endured as a conceptual framework precisely because it captures intuitions about developmental stability and canalization. Our findings refine rather than replace this metaphor (Fig. 1). The valleys are indeed deep, carved by canalization to ensure reliable development. But this same canalization flattens the ridges near valley floors, creating a landscape where mature phenotypes are poised for potential transitions despite the protective valleys that brought them there.

This revised landscape provides a unifying framework for understanding how biological systems achieve both the stability needed for reliable function and the plasticity required for adaptation, wound healing, and evolution. The attractor-centric measure introduced here offers a path toward quantifying this balance in real biological systems and predicting when and how phenotypic transitions will occur.

## Acknowledgments

The authors thank Maria Siskaki for useful comments and suggestions. MW was funded in part by the National Science Foundation, grant number DMS-2424635, and by the NIH, grant numbers R01 AI135128 and R01 HL169974-01. CK was funded in part by the Simons Foundation, grant number 712537, and the National Science Foundation, grant numbers DMS-2424632 and DMS-2451973.

## Data and Code Availability

All code underlying this study is freely available at https://github.com/ckadelka/AttractorCoherence.

## Supplementary Materials

**Figure S1:**
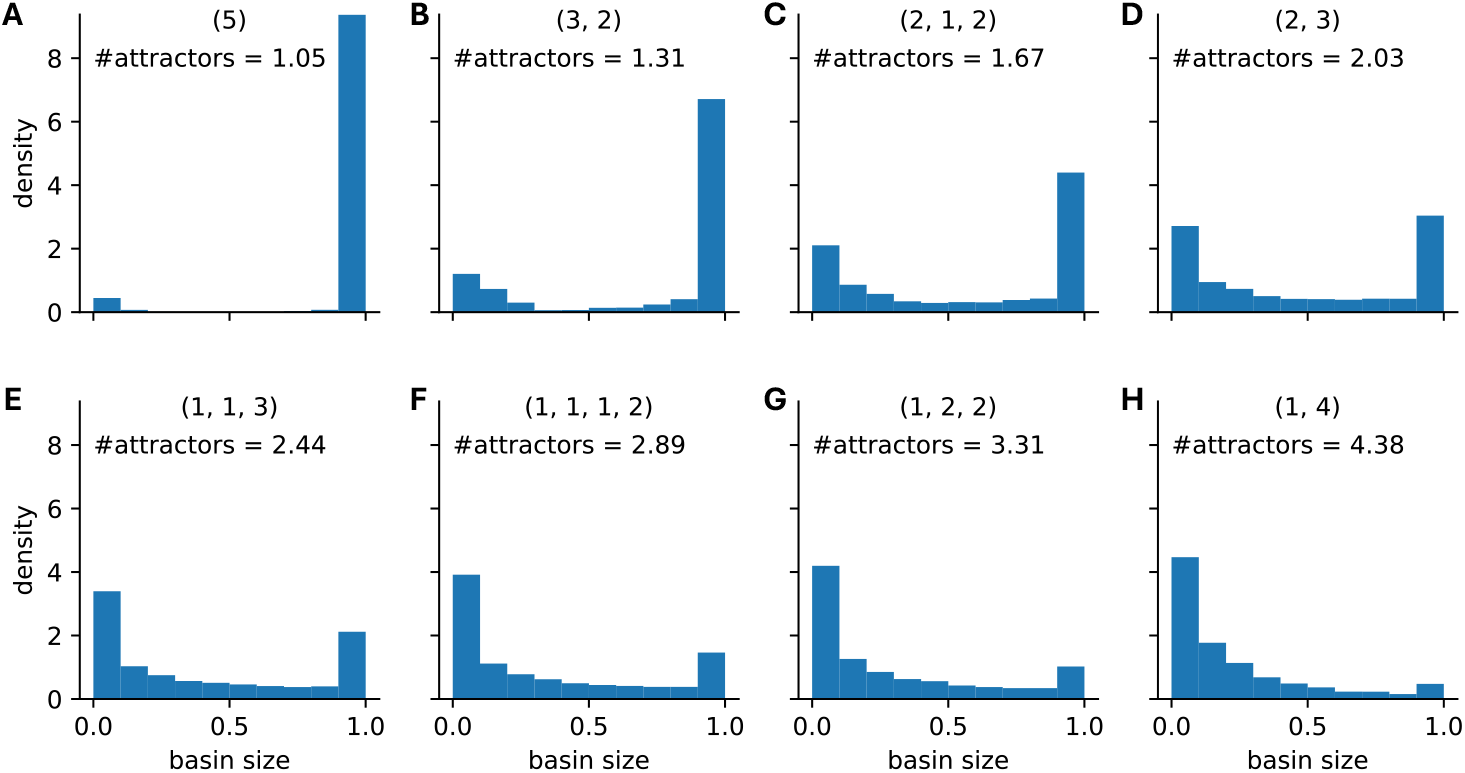
Basin size distribution for different types of nested canalizing Boolean networks. For 10,000 random 12-node BNs governed by 5-input nested canalizing update rules of different layer structure (specified above each panel), the distribution of the basin size is shown, in addition to the average number of network attractors.

**Figure S2:**
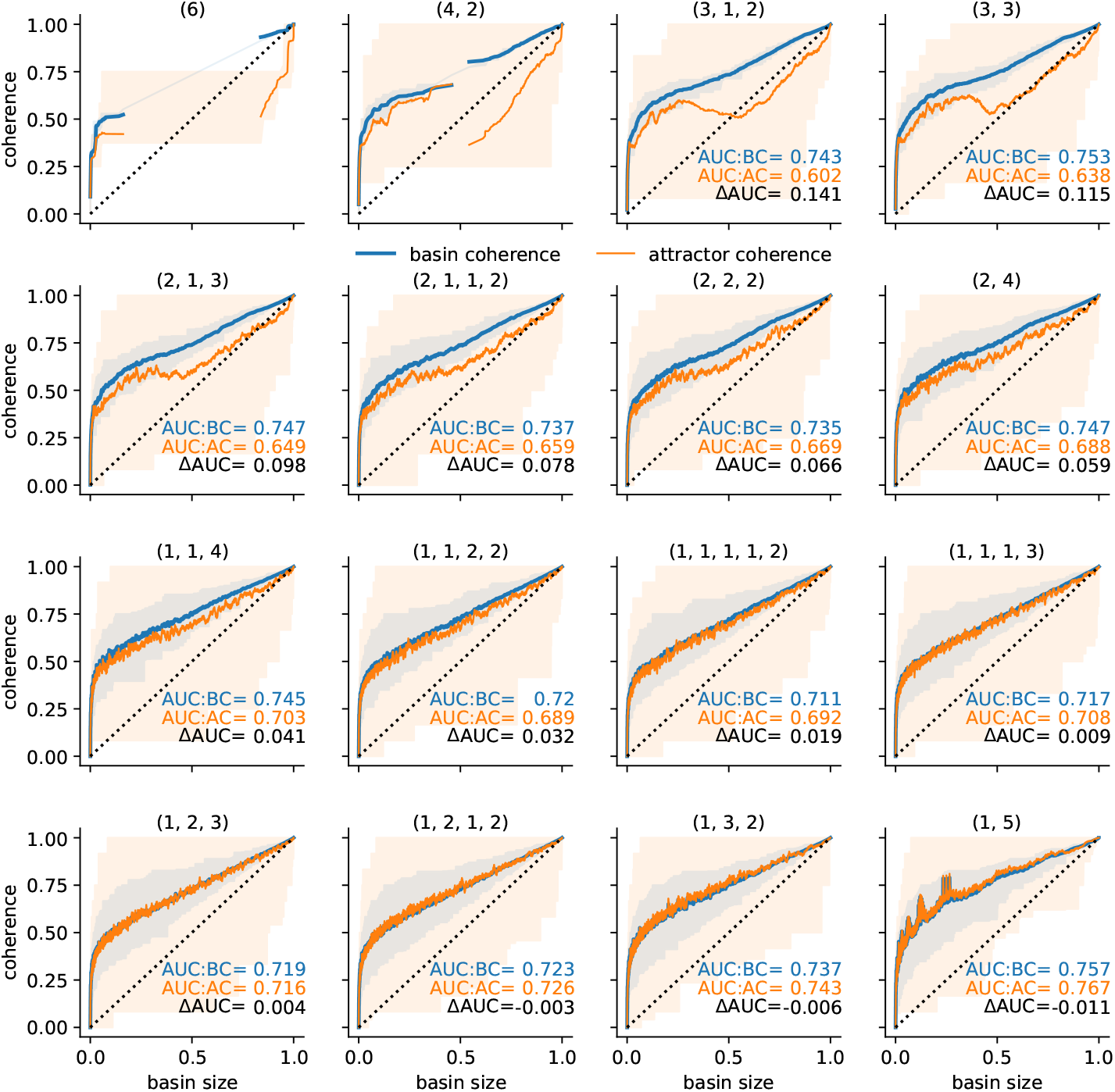
Robustness of nested canalizing Boolean network attractors and their basins. For 10,000 random 12-node BNs governed by 5-input nested canalizing update rules of different layer structure (specified above each panel), the coherence of each attractor (orange) and the coherence of its corresponding basin (blue) are stratified by basin size (x-axis). Shaded areas indicate the range of observed values. Lines depict rolling-window means with window size 50 and AUC is the area under the respective lines. The basin size distribution corresponding to each type of network is shown in Fig. S3.

**Figure S3:**
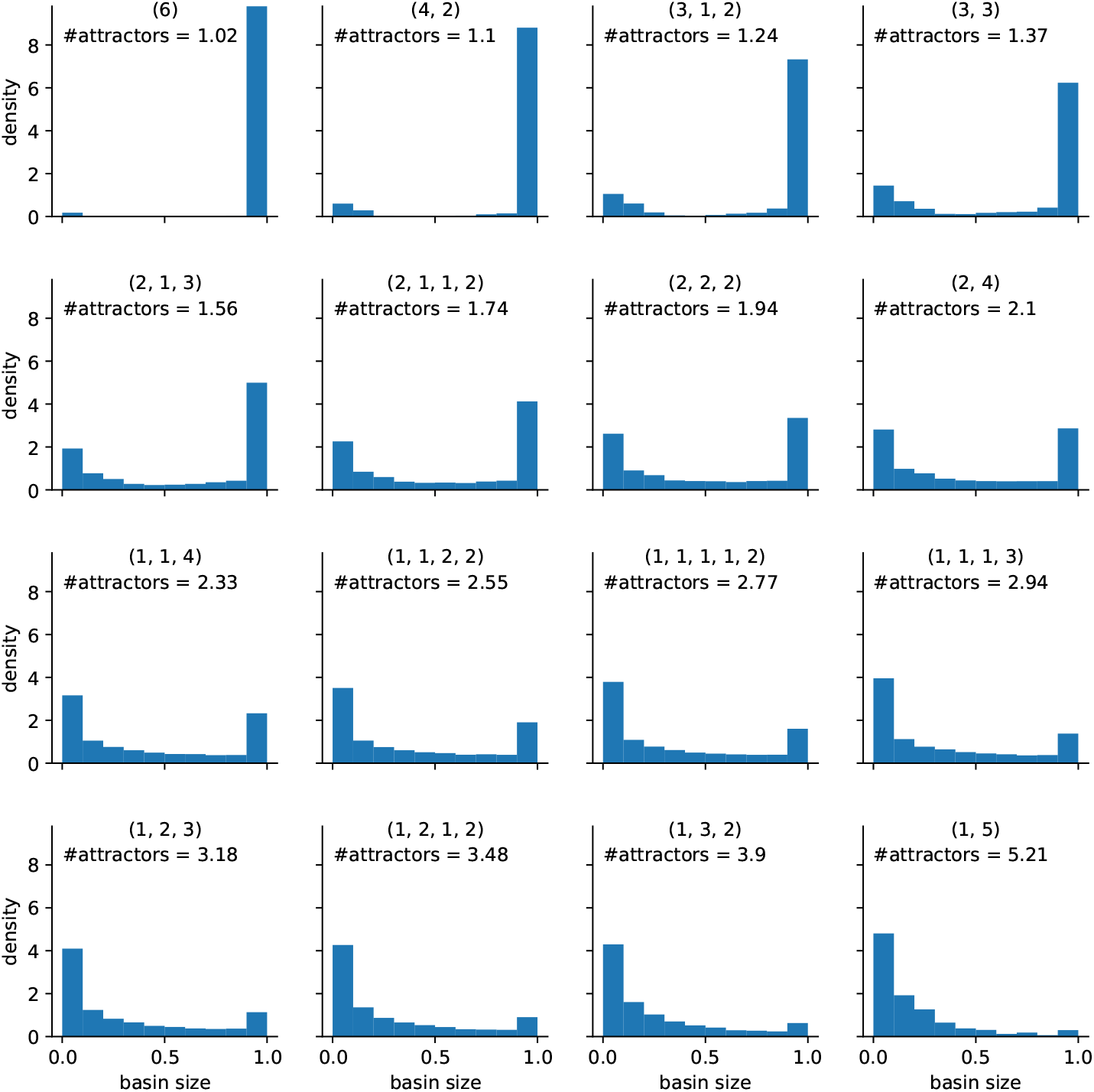
Basin size distribution for different types of nested canalizing Boolean networks. For 10,000 random 12-node BNs governed by 6-input nested canalizing update rules of different layer structure (specified above each panel), the distribution of the basin size is shown, in addition to the average number of network attractors.

**Figure S4:**
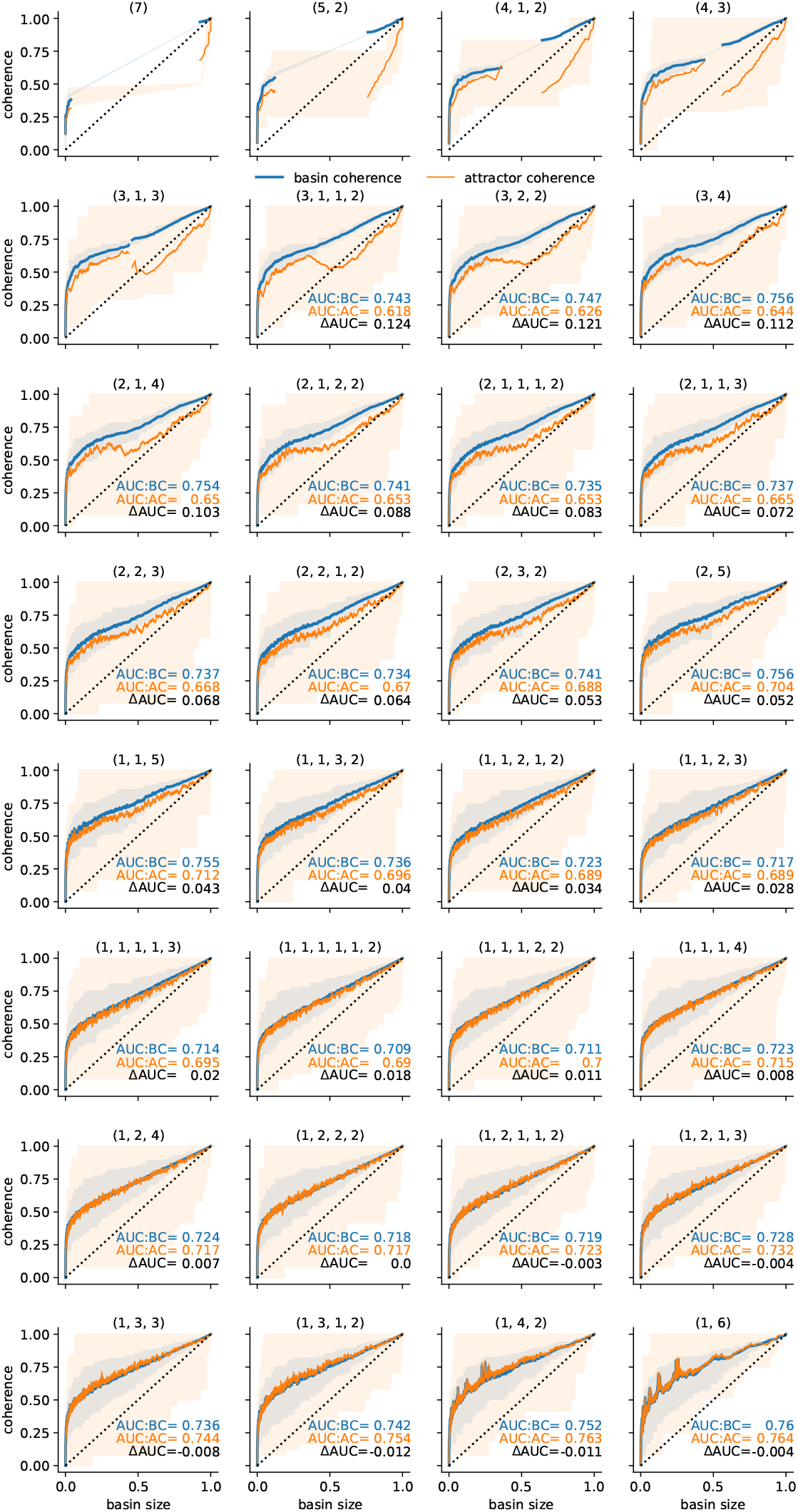
Robustness of nested canalizing Boolean network attractors and their basins. For 10,000 random 12-node BNs governed by 5-input nested canalizing update rules of different layer structure (specified above each panel), the coherence of each attractor (orange) and the coherence of its corresponding basin (blue) are stratified by basin size (x-axis). Shaded areas indicate the range of observed values. Lines depict rolling-window means with window size 50 and AUC is the area under the respective lines. The basin size distribution corresponding to each type of network is shown in Fig. S5.

**Figure S5:**
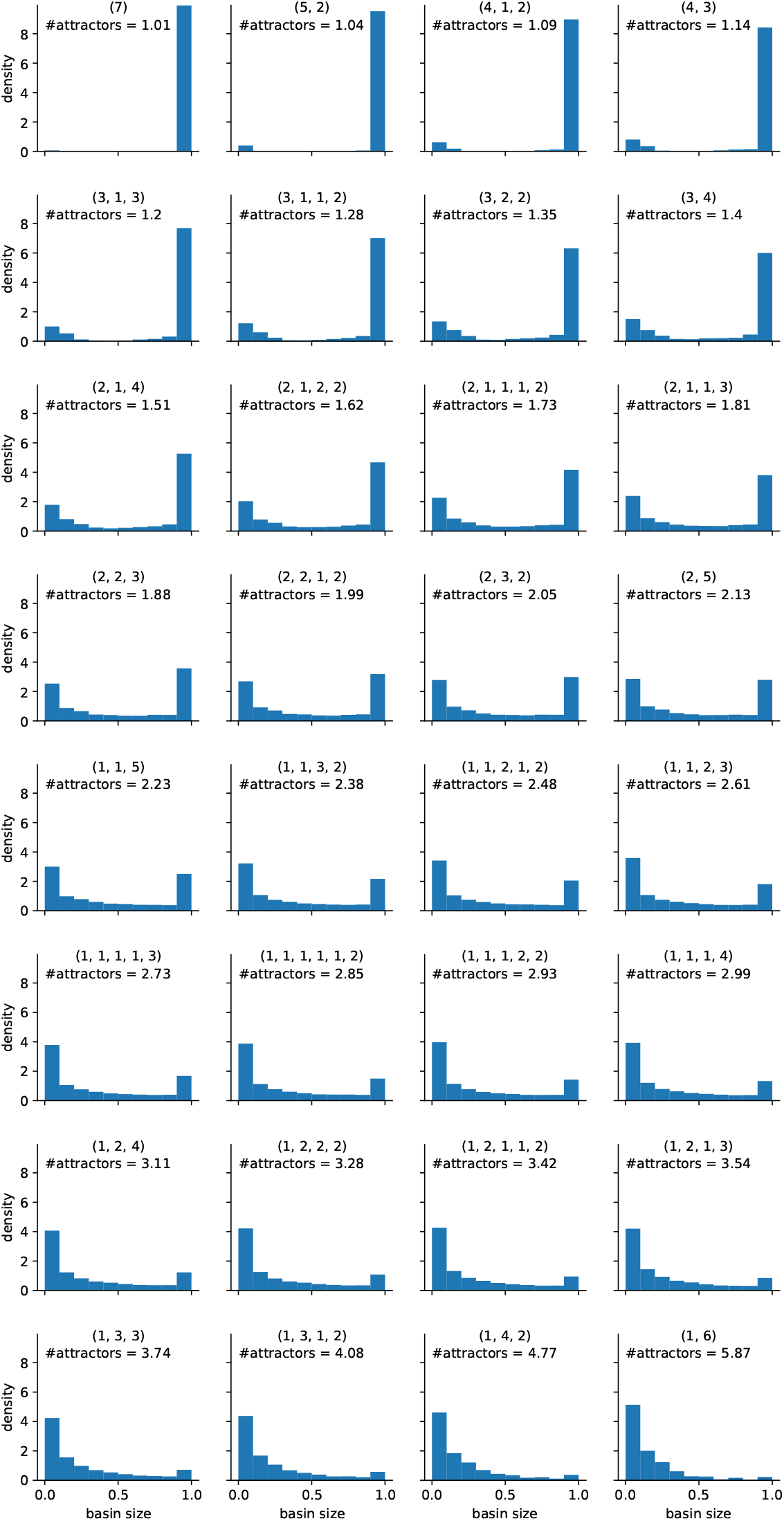
Basin size distribution for different types of nested canalizing Boolean networks. For 10,000 random 12-node BNs governed by 6-input nested canalizing update rules of different layer structure (specified above each panel), the distribution of the basin size is shown, in addition to the average number of network attractors.

